# EMhub: a web platform for data management and on-the-fly processing in scientific facilities

**DOI:** 10.1101/2024.08.12.607607

**Authors:** Jose M. De la Rosa-Trevin, Grigory Sharov, Stefan Fleischmann, Dustin Morado, John C. Bollinger, Darcie J. Miller, Daniel S. Terry, Scott C. Blanchard, Israel S. Fernandez, Marta Carroni

## Abstract

Most scientific facilities produce large amounts of heterogeneous data at a rapid pace. Managing users, instruments, reports, and invoices presents additional challenges. To address these challenges, we introduce EMhub, a web platform designed to support the daily operations and record-keeping of a scientific facility. EMhub enables easy management of user information, instruments, bookings, and projects. The application was initially developed to meet the needs of a CryoEM facility, but its functionality and adaptability have proven broad enough to be extended to other data-generating centers. The expansion of EMHub is enabled by the modular nature of its core functionalities. The application allows external processes to be connected via a REST API, automating tasks such as folder creation, user and password generation, and execution of real-time data processing pipelines. EMhub has been used for several years at the Swedish National CryoEM Facility and installed in the CryoEM center at the Structural Biology Department at St. Jude Children’s Research Hospital. A fully automated single-particle pipeline has been implemented for on-the-fly data processing and analysis. At St. Jude, the X-Ray Crystallography Center and the Single-Molecule Imaging Center have already expanded the platform to support their operational and data management workflows.

## 1. Introduction

In recent decades, the amount and complexity of data across many scientific fields have massively increased. This surge in data has paved the way for new research opportunities and innovative approaches to designing, conducting, and evaluating experiments. The biological sciences have also experienced an increase in data generation due to high-throughput instruments and data collection methods, making biological disciplines more data-driven.

Cryogenic Electron Microscopy (CryoEM) has experienced a “Resolution Revolution” thanks to recent technical breakthroughs (Kuhlbrandt, 2014*a*; Kuhlbrandt, 2014*b*; Bai *et al*., 2015). These advancements have led to the determination of larger macromolecular structures at high resolution of details. As a result, the computational requirements (hardware and software) have reached unprecedented levels. One pivotal breakthrough is the development of direct electron detectors, which have increased the speed of data acquisition and the volume of generated data (Wu *et al*., 2015). These new detectors can capture movies instead of single images, and they can cover a larger field of view. Furthermore, the high level of automation allowed the collection of tens of thousands of movies in a 24-hour microscopy session. The combination of larger formats for movie acquisition and faster detector readout has significantly increased the amount of data collected for each CryoEM processing project (Baldwin *et al*., 2017).

CryoEM facilities face the challenges of dealing not only with large volumes of data but also the complexity of data and associated metadata. Different microscopes, cameras, and acquisition software can be used, making it difficult to consistently track all ongoing experiments and their parameters. Furthermore, advancements in CryoEM have made the technique more appealing to researchers in other fields, resulting in increased demand and the establishment of many more facilities globally, following two main models (Alewijnse *et al*., 2017). The first type is similar to synchrotron facilities used by X-ray crystallographers, providing access to high-end CryoEM instruments for external users. The second type consists of core facilities in educational institutions, primarily serving internal users affiliated with the institution. There are also hybrid facilities that fall somewhere in between these two models, serving a mix of users and operational workflows.

Data management is crucial for CryoEM and other scientific facilities. However, many facilities lack proper computational infrastructure and software for their operations. Fig. 1 illustrates a typical data management workflow in CryoEM. Access to different instruments must be coordinated and scheduled based on user type, facility policies, and the data acquisition technique. Users may need multiple sessions on several instruments for the same project, and keeping track of related experiments can help with planning and timely execution. Having on-the-fly data processing workflows is also advantageous for continuously assessing sample and data quality and instrument performance with minimal delay. This helps tremendously to ensure efficient instrument usage and high-quality services for users.

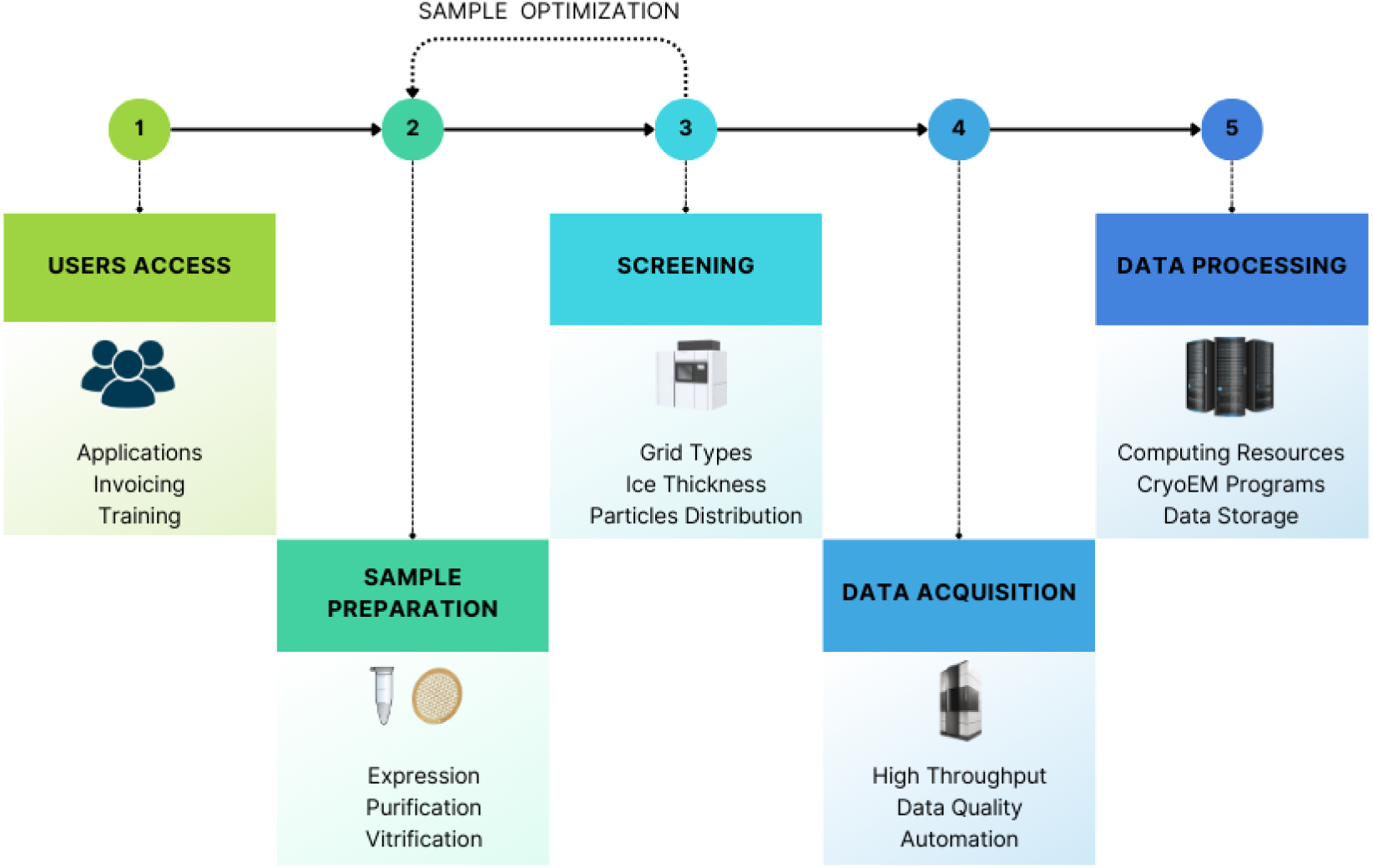
Overview of the data management workflow for a CryoEM Facility: Users request access to the facility and instrument time. Biological samples are prepared, optimized, and screened, often requiring iterations and multiple microscopy sessions. Data is acquired using high-end instruments, and some on-the-fly data processing is essential to validate its quality and the instrument’s performance.

In 2017, when the CryoEM Swedish National Facility opened, we used three different web applications and several scripts to manage users, applications, bookings, and conduct on-the-fly processing. As our facility expanded with the purchase of new instruments and an increase in number of users, the existing solution was unable to meet the growing needs. Generating annual reports and session summaries for invoicing became troublesome and time-consuming. Although we explored applications used by other CryoEM centers, none of them fit our requirements or were easy enough to customize or expand. As a result, we began developing EMhub to address our specific needs, and over time, it evolved into a more general framework to support various operations. In the following sections we will outline the different components and features of the application.

## 2. Basic Concepts and Operation

EMhub is a web framework designed to support the daily operations of a scientific facility. It includes functions for user and instrument management, project tracking, invoicing/reporting, and data transfer and processing. The system allows the creation of different types of *Users*, such as site admins, facility staff, and principal investigators (PIs). *Users* are typically organized into labs by their PI and can also be associated with *Applications* (e.g., from different grants, departments, or universities).

Staff members or site administrators can define *Resources* that can be allocated to users and may have an associated cost per *Session*. These resources can be instruments such as microscopes or services. Users will access the resources through the *Booking Calendar*, following rules set by the facility. *Booking* rules can be configured per resource and defined for different groups of users within the same lab or *Application*. *Sessions* are carried out by users via an instrument booking and typically involve data collection for a specific sample. Depending on the user’s experience, a facility manager may be assigned to manage the operation of that session.

All sessions and other research experiments can be linked via *Projects*. One user will be the project owner, while others can be added as collaborators. Within each project, different entries can be created to annotate various events, providing complete traceability and accountability for the research conducted, as well as all relevant parameters. Some types of entries allow the generation of PDF reports for the tasks performed.

### 2.1. Users and Applications

In EMhub, there are four main types of users: principal investigators (PIs), lab members, facility staff (managers), and admin/developers. PIs are independent researchers who run a lab and have privileges for lab-related operations in the application. Lab members are non-admin users associated with a specific PI, the primary unit for invoicing and reporting. They inherit their PI’s booking permissions, such as applications, booking slots, and resource allocation quota. Facility staff accounts have permission to bypass most rules and restrictions and are intended for use by facility personnel to handle cases outside normal operations. Admin/developer roles have higher-level permissions for configuring, maintaining, deploying, and troubleshooting the web application, in addition to addressing software operations issues.

*Applications* are used to organize PIs and their lab members within a logical structure. This concept can be adapted to support the allocation of access time and other services in different ways depending on the facility. In national facilities that serve multiple institutes, applications could be used to represent each external institution. This allows to establish distinct booking rules for each instrument and to maintain comprehensive statistics. In the case of an internal facility serving users from a single institute or university, applications can represent different departments or grants associated with PIs. Additionally, applications have a status property (“active” or “inactive”) that helps manage multiple active applications and create new ones over time.

### 2.2. Resources and Bookings

*Resources* are used in EMhub to represent instruments or services the facility provides to its users. Examples of instruments in a CryoEM center are microscopes, plunge freezers, and other sample preparation devices. An example of a service would be regularly scheduled drop-in sessions to offer users support with planning, preparation, or data processing for their projects.

*Resources* are central to the *Bookings* and time allocation for *Applications*. Each resource can have unique booking rules and exceptions for specific applications. For instance, we can establish minimum and/or maximum booking time, cancellation time, and session cost per day for *Resource X*. For example, a rule might be implemented for a 200 kV screening microscope requiring single-day bookings, while for a high-end 300 kV microscope, the rule could entail a minimum booking of one day and a maximum booking of two days. Furthermore, session costs could vary for each type of resource, and users may be permitted to book some resources directly while approval from facility staff might be required for others.

Specific properties are associated with each resource, which makes it easier to work with them in the system. These include the ability to define the resource name, color, icon, and tags. It is also possible to change the status of a resource from “active” to “inactive” to prevent bookings or any other operation. *Resources* can be added or removed from the *Resource List*, and the properties of each resource can also be configured. It is important to note that these actions require the user to have admin or manager roles. More detailed information about Resources can be found here: https://3dem. github.io/emdocs/emhub/user_manual/resources_bookings.html#resources *Bookings* are used to manage resource access based on the rules for each resource and the lab or application affiliation of a user. One helpful type of booking is called a “slot”, which allows access within a specific date range to certain applications (user groups) while denying access to others. This feature is particularly useful for a national facility that serves users from different universities. In this scenario, an *Application* can be defined for each university. Slots can enable users from University A to book in one week and from University B the following week. This approach allows the facility to allocate overall time efficiently while giving users the freedom to book sessions independently according to their own needs.

Only admins or managers can set slots and other particular types of bookings (e.g., downtime, maintenance). Recurring events can also be defined, allowing the setup of bookings for periodically scheduled maintenance or recurring services (e.g., drop-in sessions every Wednesday). Each booking has a designated user who is the owner. In most cases, a facility staff member will be assigned to take care of the session that day. Additionally, an *Experiment* form can be associated with the booking, allowing users to provide extra information. This can help the facility staff to plan the required work for that session more effectively. The Experiment form can be customized to reflect the facility’s workflow and needs.

The *Booking Calendar* page shows the bookings for all resources or a subset of them. The *Dashboard* page is another common place where users can view bookings for the current week and past, ongoing, or future sessions. Other pages may also have links to specific booking dialogs for displaying or editing the information. More information about bookings can be found at https://3dem.github.io/emdocs/emhub/user_manual/resources_bookings.html#bookings.

### 2.3. Sessions and Dashboards

*Sessions* represent the execution of an experiment, typically involving data collection on a specific instrument as per a scheduled booking. The necessary actions and data management for a session can vary among facilities. This may involve creating folders or user accounts and transferring and processing data. Sessions have the potential to streamline communication with facility users by providing essential information such as data collection parameters, data location, and instructions for data download on a dedicated page. From the facility’s perspective, *Sessions* allow tracking of essential metrics, such as the number of users served and the amount of data collected.

Even though bookings can be viewed from the *Calendar* page, the *Dashboard* (see Fig. 2) offers a more focused display. It helps with session planning by showing the bookings for each instrument for the current and following week. This is useful for scheduling instrument usage and planning the staff needed for each session. The EMhub web framework allows the *Dashboard* page to be customized in order to suit the needs of the facility.

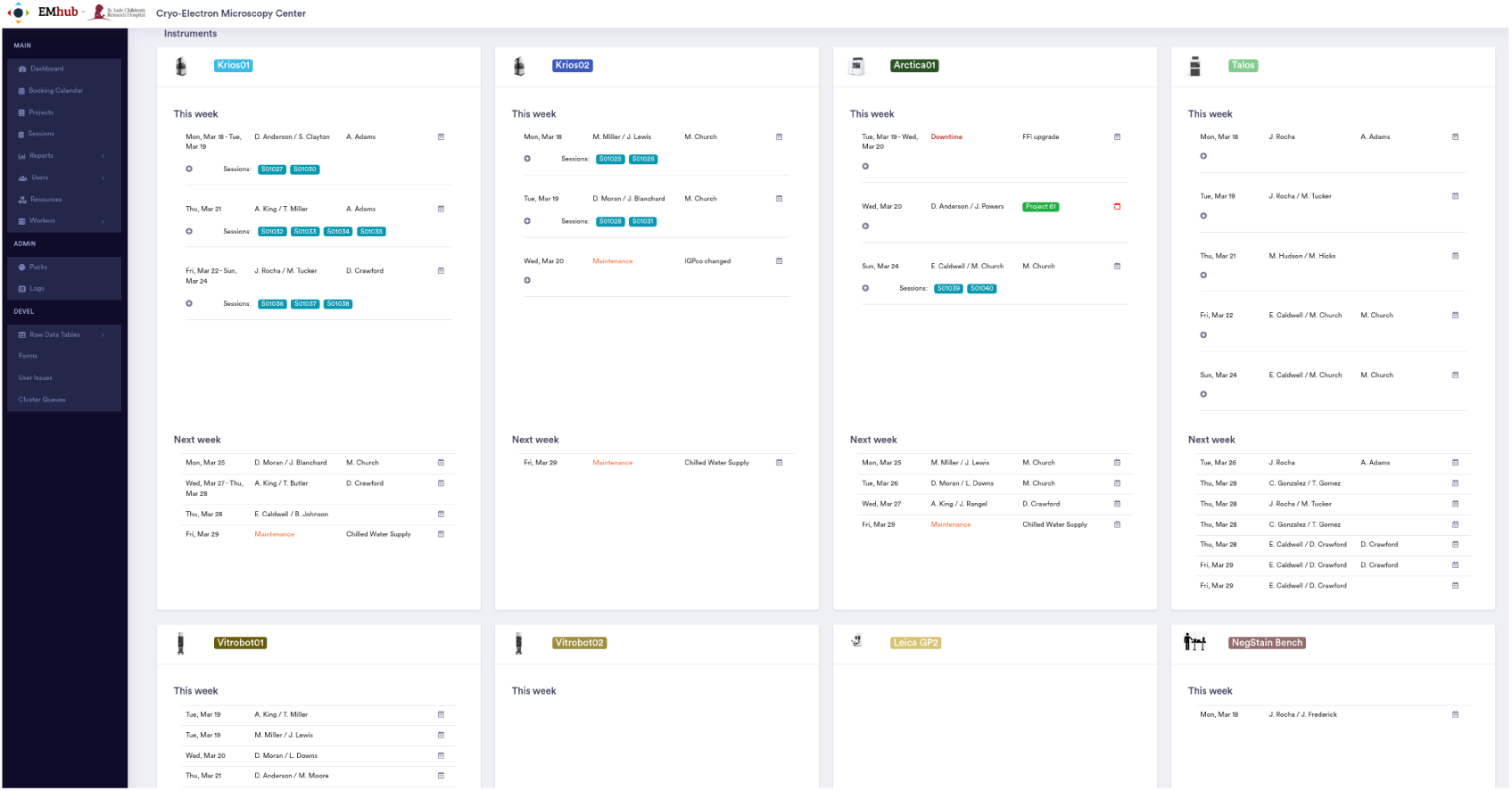
Example of the Dashboard page used at the CryoEM center in St. Jude Children’s Research Hospital. The page displays bookings for the current week and allows requests for the next week. These requests are used during the facility’s weekly meeting to schedule the next bookings and assign staff.

### 2.4. Project and Entries

A *Project* in EMhub is a valuable way to group and document events (like bookings and sessions) related to a research project over time. Users can see the entire timeline of a project from its creation. It also enables the collection of various data and metrics related to executed sessions, such as the number of days of instrument usage, images collected, and the total amount of data.

Different types of *Entries* can be added to a project timeline, providing additional annotations to a project. Using the EMhub configuration, managers can define the available entries and customize the information for each one. These entries can be used to perform various operations and later gather the information for reporting purposes. More information about projects and entries can be found at https://3dem.github.io/emdocs/emhub/user_manual/projects_entries.html.

### 2.5. Reports and Invoices

Reporting and invoicing are crucial aspects for most facilities to manage. While some institutions have a system in place, it may not track instrument usage and data collected in a comprehensive way. EMhub addresses this gap by enabling the generation of detailed reports and invoices. Having this information accessible in an organized structure allows further development of tools for analysis and reporting. These tools are essential for demonstrating the facility’s performance to evaluation committees or funding agencies. They can also aid in making informed decisions regarding current and future needs, such as additional personnel or equipment acquisitions.

EMhub includes a built-in report page showing the overall instrument usage, as depicted in Fig. 3A. Managers can choose which instruments to include in the report and specify a date range (e.g., last month, current year, previous year, etc). After updating the plots with the selected parameters, a pie chart displays the instrument usage, downtime, and maintenance. Additionally, a list by PIs and/or Applications allows the review of all bookings contributing to the period under inspection. Fig. 3B displays a session report over the selected period, showing the number of monthly sessions, the staff assignment, and the amount of data collected. The total number of projects and newly created ones can be inspected as shown in Fig. 3C. The screenshot in Fig. 3D shows the different types of entries in the timeline of a project.

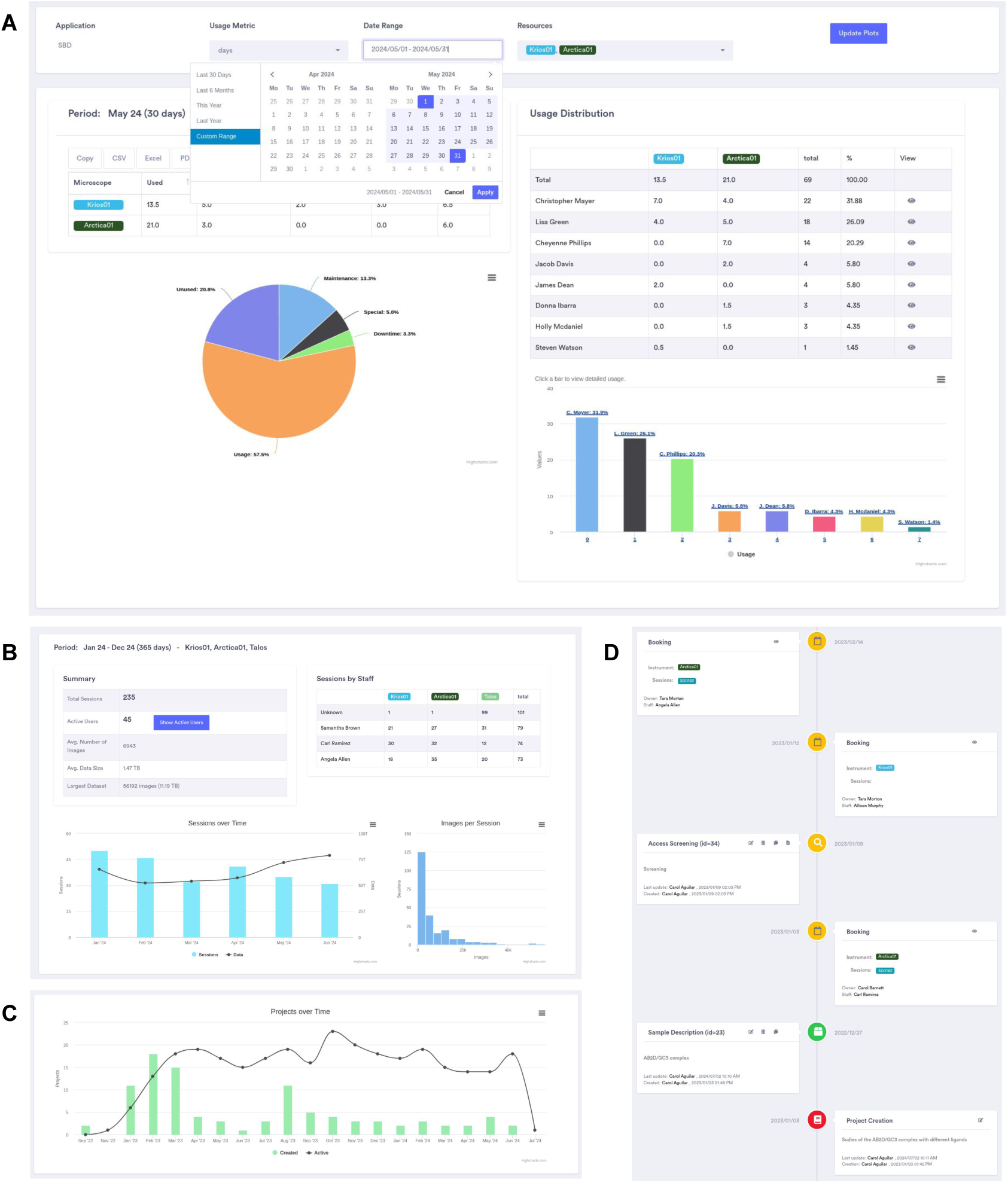
Different reports generated in EMhub. A) Instruments usage report for a selected period, grouped by labs. B) Overall report for all sessions, staff assignments, and data collected. C) Total number of projects and new ones over time. D) A project’s timeline showing different types of entries.

## 3. Use Cases, Extensions, and Software

### 3.1. EMhub at the Swedish National CryoEM Facility

The Swedish National CryoEM Facility provides access to cutting-edge equipment and expertise in single particle CryoEM, cryo-tomography (CryoET), and MicroED through two nodes belonging to SciLifeLab, one in Stockholm and one in Umeå. At the SciLifeLab node in Stockholm, researchers can perform single-particle CryoEM, MicroED, and CryoET for plunge-frozen samples, utilizing a Talos Arctica for sample optimization and two Titan Krios for high-resolution data collection. The node in Stockholm also offers remote drop-in service for image processing support. The Facility accepts two types of applications: Rapid Access (RA) and Block Allocation Group (BAG). RA applications can be submitted at any time and are evaluated for technical feasibility within the allocated time in the following quarter. BAG applications are evaluated annually by a national Project Evaluation Committee based on scientific merit and technical feasibility.

In the Stockholm node, EMhub has been in use since January 2019. Before that, several software programs were used for data management, like in many other facilities. Initially, there was a web portal where users and PIs registered and submitted applications for instrument access. There was also a separate booking calendar with a different user base unrelated to the web portal. Some custom scripts were also used to create folders for data collection and perform on-the-fly data processing. Furthermore, a significant amount of communication with users occurred via email to provide the necessary details for a session (such as acquisition parameters and data transfer credentials), and there was an urgent need to document the progress of projects across different sessions.

EMhub was developed to provide a practical solution for all of these needs. It allowed interaction with the Portal to import users and applications while centralizing all information under a single system. Access to instruments, bookings, and sessions are all managed through EMhub. When a session is created, a corresponding folder and the credentials for data download are automatically generated (based on the user’s application and lab). Users can access all this information on the page of a Session. The facility typically allocates time among different applications using slots in the calendar. Users from a specific application can book within the allocated slot days, ensuring fair resource utilization and flexibility for experiment planning. EMhub also provides drop-in services for project support, which can be booked on specified days. The facility staff extensively uses projects to document the progress of ongoing collaborations. Additionally, numerous entries related to sample preparation, screening, and collection enable tracking of the position of sample grids in the storage dewars. This allows facility staff to record the screening process for specific grids, and another person can easily access the information during the data collection session. It also provides research groups with better overall management of the sample optimization process and enables a smooth transfer of project responsibility if necessary.

Another feature heavily used by the node in Stockholm is the generation of invoices and reports. EMhub allows the definition of different invoice periods (e.g., quarterly in a year) and accounts for the sessions in that period for each lab. The invoicing information is more transparent for the PIs, who can review all the sessions per user in that period and the associated costs. At the end of a period, the facility generates an invoice report that is handed to the financial administrators for actual invoicing. Reports are also crucial for presenting yearly facility numbers and performance for evaluation purposes.

### 3.2. EMhub at the CryoEM center in St.Jude Children’s Research Hospital

The CryoEM Center at the Department of Structural Biology in St. Jude Children’s Research Hospital provides state-of-the-art instruments and experienced staff scientists to support internal research. The center enables St. Jude researchers to explore intricate biological structures, such as large macromolecular complexes at almost atomic resolution. A key objective is to offer comprehensive support and training to investigators at all levels across the entire single-particle workflow, from sample preparation to data collection, image processing, 3D reconstruction, and modeling. The CryoEM Center is equipped with a 300 kV Titan Krios and a 200 kV Talos Arctica, both fitted with a K3 direct electron detector and a BioQuantum energy filter. The recent installation of another Titan Krios with a Falcon 4 detector and SelectrisX energy filter has also been completed. The Center also houses advanced sample preparation, screening, and optimization equipment.

Despite not being a national facility, the increase in equipment and the recruitment of new PIs and lab members pose some data management challenges to the center. After some modifications and extensions, EMhub was deployed for daily usage at the center in March 2023. Previously, instrument access was requested via email, with experiment information in a Word document, which was often redundant or inaccurate. All these processes were moved into EMhub, taking advantage of the flexibility of the Entries in a Project. A new type of entry (Microscope Access Request) allowed users to request a microscope, specify a desired day, and provide information about the sample and help needed for the session. A new dashboard page was developed to show all requests for the next week, which could be converted into bookings, and center staff could be assigned. Users can review the assignment after the weekly time allocation and plan accordingly.

Projects have also helped track the usage of instruments over time in each lab, as well as other statistics such as the number of images/data generated per session and overall in a project. Similarly, reports enable the center to monitor facility operations, especially the usage of microscopes (e.g., time distribution, downtime, maintenance, etc).

### 3.3. Data transfer and On-The-Fly Processing

When a data collection session started at the CryoEM center at St. Jude, some scripts transferred the data to a buffer server and then to the centralized shared filesystem. However, there was no explicit link between the raw images and the booking, which made it challenging to collect overall statistics. With EMhub in place, it became easier to relate the current data collection with the booking user and their corresponding PI. The transfer script was integrated with EMhub, and the information on transferred files was made available via the session’s page.

For the on-the-fly processing, the user manually launched a Relion pipeline (Fernandez-Leiro & Scheres, 2017). The pipeline performed motion correction, contrast transfer function (CTF) estimation (Rohou & Grigorieff, 2015), blob picking, and 2D classification (Scheres, 2012; Scheres, 2015; Kimanius *et al*., 2016). Additionally, facility staff generated and used some plots to monitor data quality during acquisition. When EMhub was introduced, one of the initial tasks was to integrate booking/session information with the processing pipeline. The existing plots were adapted for the web, and a new Session Live page was created to monitor data collection progress and processing results, as shown in Fig. 4. Some input parameters were retrieved from the microscope and data collection settings, eliminating the need to re-enter them into the pipeline.

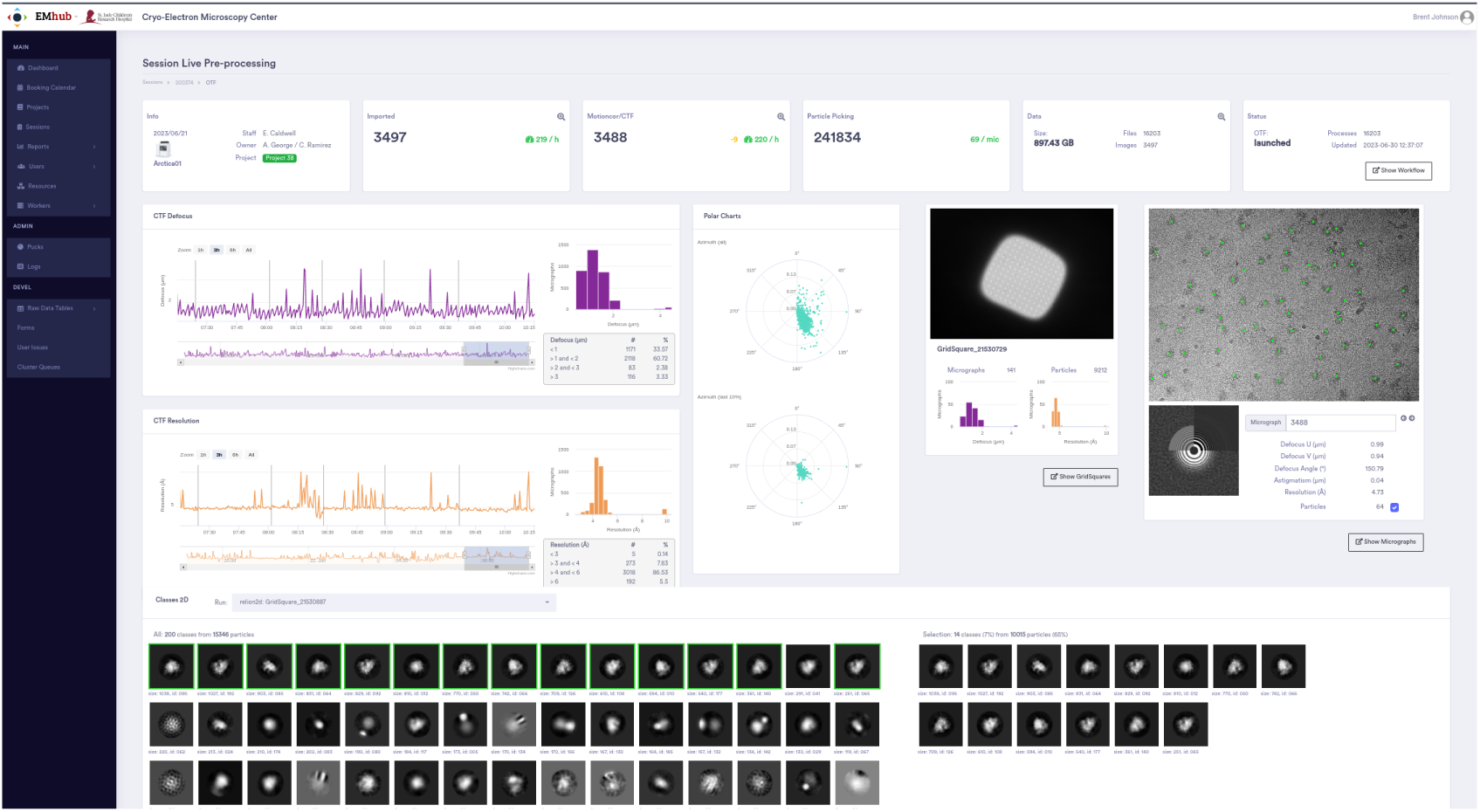
EMhub Page to monitor data collection and on-the-fly processing.

To improve our on-the-fly capabilities, we have implemented support for Scipion (de la Rosa-Trevín *et al*., 2016) workflows as the default pipeline. We have made some modifications to the general processing steps. First, we sped up the motion correction step using Motioncor2 (Zheng *et al*., 2017) in batch mode. Then, we changed the picking process by using the general trained model from Cryolo (Wagner *et al*., 2019), which has proven to work well for most samples and provides a particle size estimate. This has allowed us to automate the pipeline without any human intervention. After obtaining the particle size from Cryolo, we have adjusted the box size for extraction and 2D classification accordingly. Another change is the scheduling of 2D classification in batches, where each batch consists of particles extracted from the micrographs of a specific grid square. Upon completing the 2D classification job, we conduct a 2D class selection to gain a rough idea of “good classes” and the underlying “good particles”. This approach ensures that a similar number of particles are used as input for each 2D classification job, resulting in quicker completion and providing more realtime feedback about the sample quality. Additionally, we can now gather statistics about the number of good particles per grid square and CTF metrics such as defocus, astigmatism, and resolution.

### 3.4. EMhub for non-CryoEM Facilities

#### 3.4.1. X-ray Center

The Biomolecular X-Ray Crystallography Center (XRC) in the St. Jude department of Structural Biology provides diverse St. Jude researchers access to tools, facilities, and professional assistance for all aspects of high-resolution biomolecular structure determination by X-ray crystallography. Center staff and trained walk-up users employ robotic instruments for efficient screening of large numbers of crystallization conditions for both soluble and membrane proteins. Along with a host of other instruments, protein crystallization imagers are available for automatic image capture and drop reporting. Once crystals are obtained, they can be assessed via an in-house X-ray diffraction system, and the facility manages external relationships with several synchrotron beamlines affording St. Jude researchers regular access to the most advanced light sources and cutting-edge technology.

The XRC has utilized a variety of means to manage access to its on-site resources and the synchrotron time it stewards. The most recent approach before EMHub involved customized calendars implemented in Microsoft SharePoint, incorporating a variety of custom fields and business rules. Even with those aids, however, the XRC staff spent an undue amount of time coordinating synchrotron time requests and scheduling, a lot of that via e-mail communication, spreadsheets, and other documents. Not only was the information that needed to be gathered and collated too diverse to be easily handled via SharePoint, but XRC procedures involve a multi-step, back and forth workflow to allocate shared beam time for samples, and this wasn’t easily modeled or automated in SharePoint.

In addition to serving well straight out of the box for scheduling access to the XRC’s on-site instruments, EMHub proved flexible and extensible enough to support the whole beamtime request and approval workflow. Through EMHub, users now provide the needed information about the experiments they want to perform; XRC staff vet those and allocate available time; users provide essential information about their harvested crystals and the puck in which they are stored; and center staff prepare detailed schedules using a simple drag and drop utility and automatically prepare time-sensitive beamline-required documents. Furthermore, all this information persists in one place – EMHub’s database – that can be referenced later to resolve questions ranging from data provenance to distribution of resources. Additionally, XRC staff have found the EMHub REST API, and especially the ability to define custom endpoints, to be useful for automating post hoc data management procedures. This includes efficient delivery of synchrotron data to institutional storage according to PIs.

#### 3.4.2. Single-Molecule Imaging Center

The mission of the Single-Molecule Imaging Center (SMC) at St Jude is to make single-molecule imaging methods broadly accessible to scientists at St. Jude and beyond, including those without extensive technical background in microscopy and biophysics. The SMC houses state-of-the-art, custom-built instrumentation for single-molecule fluorescence imaging, including confocal time-correlated single-photon counting (TCSPC) and total internal reflection fluorescence (TIRF) microscopes, as well as ensemble spectrometers, data analysis workstations, and other resources. Much like CryoEM, single-molecule instruments are expensive to build and maintain. An electronic booking system is essential to ensure judicious use of these valuable resources. Imaging is conducted in custom microfluidic imaging chambers fabricated with careful surface passivisation to avoid non-specific surface interactions and fluorescent contaminants. These chambers are expensive to fabricate, and thus, tracking their availability and performance is essential.

Before EMhub, instrument reservations were managed with a collection of Microsoft Outlook calendars shared with authorized users. This approach was problematic due to the lack of any mechanism to prevent scheduling conflicts, excessively long sessions, making reservations during scheduled downtime, or users accidentally modifying other’s bookings. Tracking of microfluidic chambers was handled in a separate Excel document, which suffered from similar issues and also had no direct connection with the booking calendars. EMhub presents a significant improvement on these systems by providing a centralized interface to track instrument bookings and microfluidic chambers that can enforce scheduling rules and associate all this information together with a project timeline.

Although not currently implemented, EMhub’s REST API capabilities will be utilized to track standard statistics that measure the health of the instruments and the proper execution of imaging experiments. These include signal-to-noise ratios, particle density, image sharpness (focus), illumination uniformity, optical aberrations, correction factors, etc. These statistics could be used to determine the timeline of any failures and also as a tool for monitoring users who may need additional guidance or training.

### 3.5. Implementation and Software Availability

EMhub is developed in Python (https://www.python.org/) using the Flask (https://flask.palletsprojects.com/) web micro-framework, which is known for its minimalist design and numerous extension plugins. For data persistence, EMhub utilizes SQLite (https://www.sqlite.org/) as the database back-end through the SqlAlchemy (https://www.sqlalchemy.org/) mapping layer. This provides flexibility for data operations and simplifies potential porting to a different database in the future. A key feature of EMhub is its REST API, which allows other systems to interact with the framework. This REST API has been used to extend EMhub capabilities by developing worker scripts that can run independently from the web server to perform tasks such as data transfer or on-the-fly processing. Fig. 5A shows an overview of the EMhub architecture. Fig. 5B highlights all the core components that can be extended by providing an *extra* folder inside the EMhub instance directory.

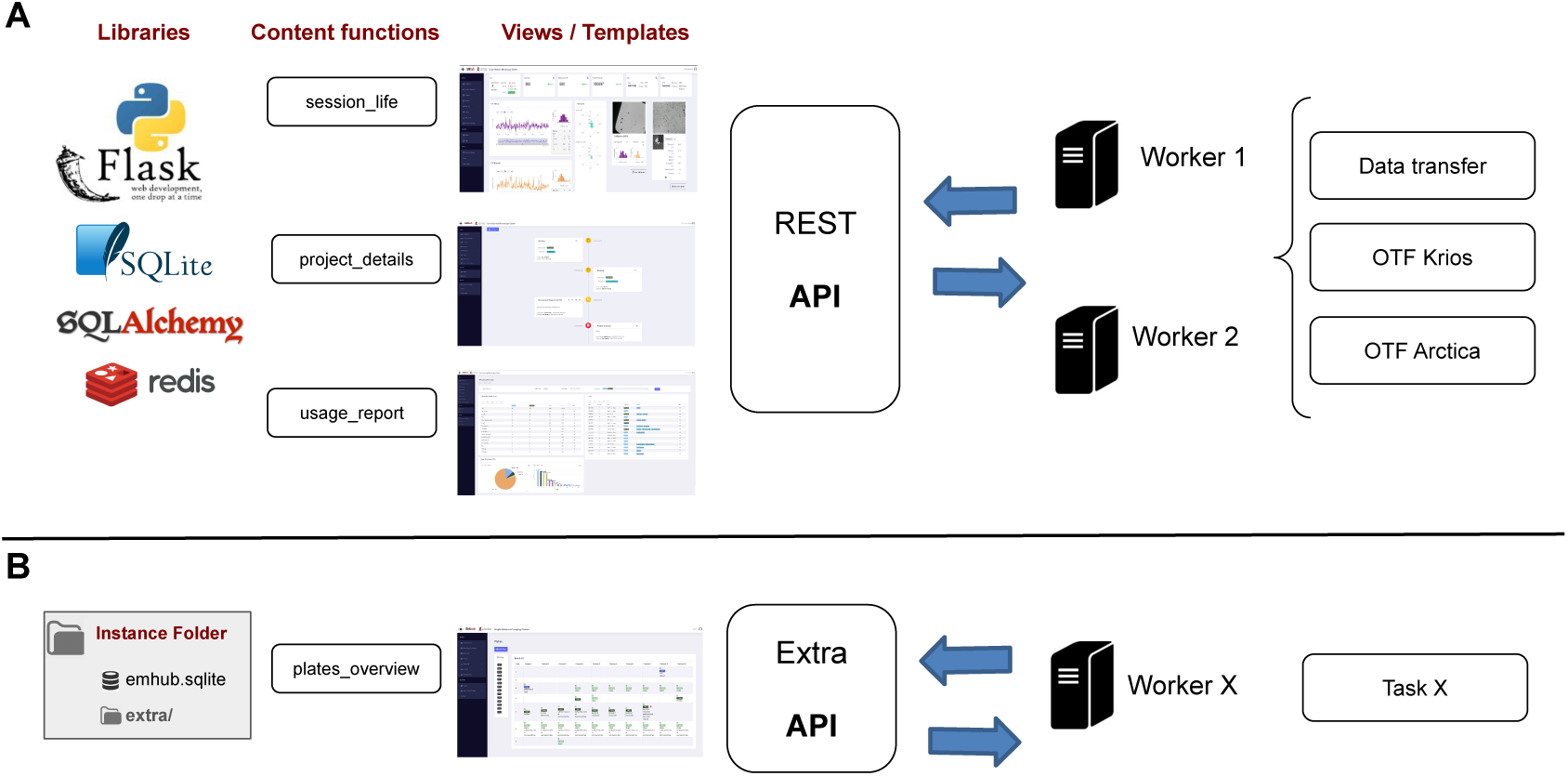
Overview of the EMhub architecture and its customization possibilities. A) The web application is based on Flask, and it is organized around template/view pages supported by content functions. A REST API allows one to write worker scripts to communicate with the application and execute other tasks. B) Views, content functions, and the REST API can be customized and extended by providing an extra folder.

EMhub is publicly available under the GPLv3.0 open-source license. Source code can be found at: https://github.com/3dem/emhub, together with extensive documentation at https://3dem.github.io/emdocs/emhub.

## 4. Discussion

This work introduces EMhub, a web framework designed for data management and on-the-fly data processing. The development of EMhub was driven by the challenges faced in the day-to-day operations of a CryoEM facility and the absence of a simple solution to address these needs. EMhub has been created to focus on simplicity while supporting tasks such as user, instrument, and booking management. Additionally, it offers comprehensive project and parameter tracking and the generation of reports and invoices. EMhub has already been successfully utilized by the Swedish National CryoEM facility at SciLifeLab in Stockholm and the CryoEM center of the Structural Biology Department at St. Jude Children’s Research Hospital. While having some needs in common, these two facilities have different user types and operational workflows, demonstrating the adaptability of EMhub in diverse scenarios.

The EMhub architecture relies on its REST API, which enables well-defined communication with the system and allows other machines to handle tasks such as account setup, data transfer, and on-the-fly processing. The core framework is highly customizable, making it possible to override specific functions or pages and to add new features. For instance, the X-ray and Single-Molecule centers have expanded the core EMhub framework to suit their particular needs.

The current implementation of EMhub has already proven valuable, and we expect it to be appealing to other facilities as well. Since different facilities have diverse needs and operations, we do not think a single monolithic solution can be ideal for every case. Keeping this in mind, we have tried our best to create a framework with many useful built-in features that can be expanded and customized. In addition, we have provided comprehensive documentation at https://3dem.github.io/emdocs/emhub/user_manual/projects_entries.html, covering topics from various perspectives (users, facility managers, sysadmins, and developers). We anticipate that EMhub will continue to evolve to meet future challenges and in response to feedback and contributions from the community.

## Acknowledgements

This work has used data generated by the CryoEM Swedish National Facility, funded by the Knut and Alice Wallenberg, Family Erling Persson and Kempe Foundations, SciLifeLab, Stockholm University, and Umeå University. The work at St. Jude Children’s Research Hospital was supported by funding from ALSAC.

The authors would like to thank all the facility staff of the CryoEM facility at SciLifeLab in Stockholm and the CryoEM center at St. Jude Children’s Research Hospital for their valuable feedback and discussions. We thank the engagement of the Biomolecular X-Ray Crystallography Center and the Single-Molecule Imaging Center at St. Jude. We are also very thankful for the tremendous support from the HPC, Cloud, and Networking teams from the Information Services Department at St.Jude Children’s Research Hospital.

## Synopsis

This article presents EMhub, a web platform designed to support the daily operations of a scientific facility. EMhub enables easy management of users, instruments, bookings, and project tracking. The application was initially developed to meet the needs of a CryoEM facility, but the versatility of its design makes it suitable to be extended to other areas.

